# Environmental DNA Reveals the Geographic Distributions of Two Eel Species, *Anguilla japonica* and *A. marmorata*, in Western Kyushu, Japan

**DOI:** 10.1101/2023.10.24.563875

**Authors:** Yurika Ono, Shimpei Tsuchida, Katsuya Hirasaka, Taijun Myosho, Shingo Fujimoto, Kenichi Shimizu, Mitsuharu Yagi

**Affiliations:** Graduate School of Fisheries and Environmental Sciences, Nagasaki University, Nagasaki 852-8521, Japan; Graduate Division of Nutritional and Environmental Sciences, University of Shizuoka, Shizuoka 422-8526, Japan; Tropical Biosphere Research Center, University of the Ryukyus, Okinawa 903-0213, Japan

**Keywords:** dispersal, eDNA, giant mottled eel, Japanese eel, river

## Abstract

Anguillid eels migrate thousands of kilometres from their spawning grounds, dispersing across a vast geographic area to fresh and brackish water habitats, where they settle and grow. Japanese eels (*Anguilla japonica*) and giant mottled eels (*A. marmorata*) are both found in Japan, although their distributions differ. However, details of these differences are unknown. We hypothesised that distribution patterns of Japanese and giant mottled eels must be different between and within rivers along the northwest coast of Kyushu, Japan. Environmental DNA (eDNA) analysis was conducted at 87 sites in 23 rivers. Japanese eel eDNA was detected in 19 rivers (82.6%) and that of giant mottled eels was detected in 8 (34.8%). eDNA for Japanese eels was detected at 6 of 9 sites in the North (66.7%), 13 of 23 sites in Omura (56.5%) and 37 of 55 sites in the South (67.3%). In contrast, giant mottled eel eDNA was detected at 1 of 9 sites in the North (11.1%), no sites in Omura and 15 of 55 sites in the South (27.3%). There was no correlation between eDNA concentrations of the two species at 10 sites in the five rivers where eDNA of both species was detected, implying that their habitat preference differ. This partially reveals dispersal and settlement mechanisms of these eel species.

## INTRODUCTION

Anguillid eels migrate thousands of kilometres from their spawning grounds, dispersing across a vast geographic area to fresh and brackish water habitats, where they settle and grow. The genus *Anguilla* contains 19 species worldwide, including three subspecies, and is widely distributed throughout tropical and temperate zones (Watanabe et al., 2004, 2009; Kuroki, 2018). Eels are typical catadromous migratory fish (Tsukamoto, 1990; Tsukamoto, 1992). This means that adult fish migrate from their nursery grounds in rivers to spawning grounds in the open ocean, and leaf-like eel larvae (leptocephali) are transported as far as 3000 km (Tsukamoto et al., 2002; Kuroki, 2018; Kasai et al., 2021). Leptocephali are transparent and laterally compressed (Kuroki 2014, 2018), which is probably adaptive for long-distance dispersal on ocean currents, with little energetic cost.

Japanese eels (*Anguilla japonica*) and giant mottled eels (*A. marmorata*), populations of which are found in Japan, although their distributions differ. In temperate regions in Taiwan, eastern China, Korea and Japan, Japanese eels are dominant in most rivers, which are typically relatively large and slow moving (Kaifu et al., 2010; Shuai et al., 2021; Tzeng et al., 1995). On the other hand, giant mottled eels are dominant in subtropical islands where relatively small rivers with fast flows are common (Itakura et al., 2020b; Kano et al., 2017; Kumai et al., 2020). They also occur occasionally in rivers in Shikoku and Kyushu, Japan, adjacent to warm currents (Ishibashi et al., 2001; Kasai et al., 2021; Tabeta, 1994). Both species are declining due to environmental degradation in habitats such as habitat loss (Chen et al., 2014; Tatsukawa, 2003) and pollution (Tabeta, 1994; Tabeta et al., 1995), fluctuations in oceanic conditions and overfishing (Jacoby et al., 2015; Kim et al., 2007; Kimura et al., 1994). The Japanese eel is listed as an Endangered (EN) species on the IUCN Red List of Threatened Species (IUCN 2022). The giant mottled eel is considered a national monument in various places in Japan, but the biology of this species is very poorly known (Mizuno and Nagasawa, 2010), and differences in distributions of the two species are thought to be due to supply by marine transport and the post-riverine environment, though such assumptions are poorly documented.

Environmental DNA (eDNA) surveys are good for revealing habitat distributions of organisms that hide in rocks and sediments, such as eels. Traditionally, sampling of eels has involved direct capture by trapping, or netting with electroshocking, which requires significant effort and is often not effective in deep or saltwater areas (Bohlin et al., 1989; Itakura et al., 2019). eDNA analysis detects DNA in water from feces and mucus, determining which species are present in an area under study (Ficetola et al., 2008; Kasai et al., 2021). A weak positive relationship between eDNA concentration and the abundance of Japanese eels suggests that eDNA can be used to estimate not only presence, but also abundance of eels in rivers (Itakura et al., 2019). Using this technique, Kasai et al. (2021) showed that Japanese eel eDNA was highly concentrated in rivers on the Pacific side of Honshu, west of the Kanto region, in the Seto Inland Sea and along the west coast of Kyushu, Japan. Japanese eel and giant mottled eel eDNA concentrations were also reported to be correlated with distances from river mouths (Itakura et al., 2019). Ono et al. (2023) also recently reported that eDNA concentrations of Japanese eel were higher at downstream than upstream sites.

eDNA surveys of eels on the west coast of Kyushu, Japan may provide insights into dispersal and settlement mechanisms of eels. Nagasaki Prefecture, located at the western end of the Japanese archipelago, is exposed to both the Tsushima Current, one of the largest warm ocean currents in the world, and the Kuroshio Current, which diverges in southwest Kyushu. It is also bordered by Omura Bay, an extremely enclosed body of water, and the Ariake Sea, an inland sea. The geographic connectivity of these marine areas and rivers is expected to influence settlement and distribution of eels. In addition, no known releases of eels have occurred in Nagasaki Prefecture (Fisheries Census by the Ministry of Agriculture, Forestry and Fisheries (2008, 2013)), implying that distributions of eels in the Nagasaki region may reflect their original distributions. Furthermore, interspecific habitat segregation occurs between these two species, probably due to species-specific habitat preferences (Matsushige et al., 2022; Shiao et al., 2003). Thus, we hypothesised that the distribution patterns of Japanese eels and giant eels would differ both within and between rivers, despite the narrow range of eels along the west coast of Kyushu.

## MATERIALS AND METHODS

### Study site and water sampling

eDNA analysis was conducted at 87 sites in 23 rivers in western Kyushu, Japan, from July to September 2022 (Fig. 1). Sampling sites (2 to 7 per river), spanned upstream to downstream habitats (Fig. S1). To avoid reduced detection probability of target species (Deniner et al., 2015; Li et al., 2018) and reduced detection of eDNA concentrations due to PCR inhibition in turbid water (Mckee et al., 2015), sampling was not conducted during increased flows due to heavy rains. The study also focused on yellow eels, which are relatively sedentary. Sampling was done during the summer months, avoiding autumn and winter months when glass and silverside eels migrate (Sudo et al., 2017). At sampling sites, water temperature, dissolved oxygen, salinity, and depth were measured using a multi-purpose digital water quality meter (YSI ProDSS, Xylem Inc., USA). In addition, flow rates, elevation and river width were recorded.

**Fig. 1.**
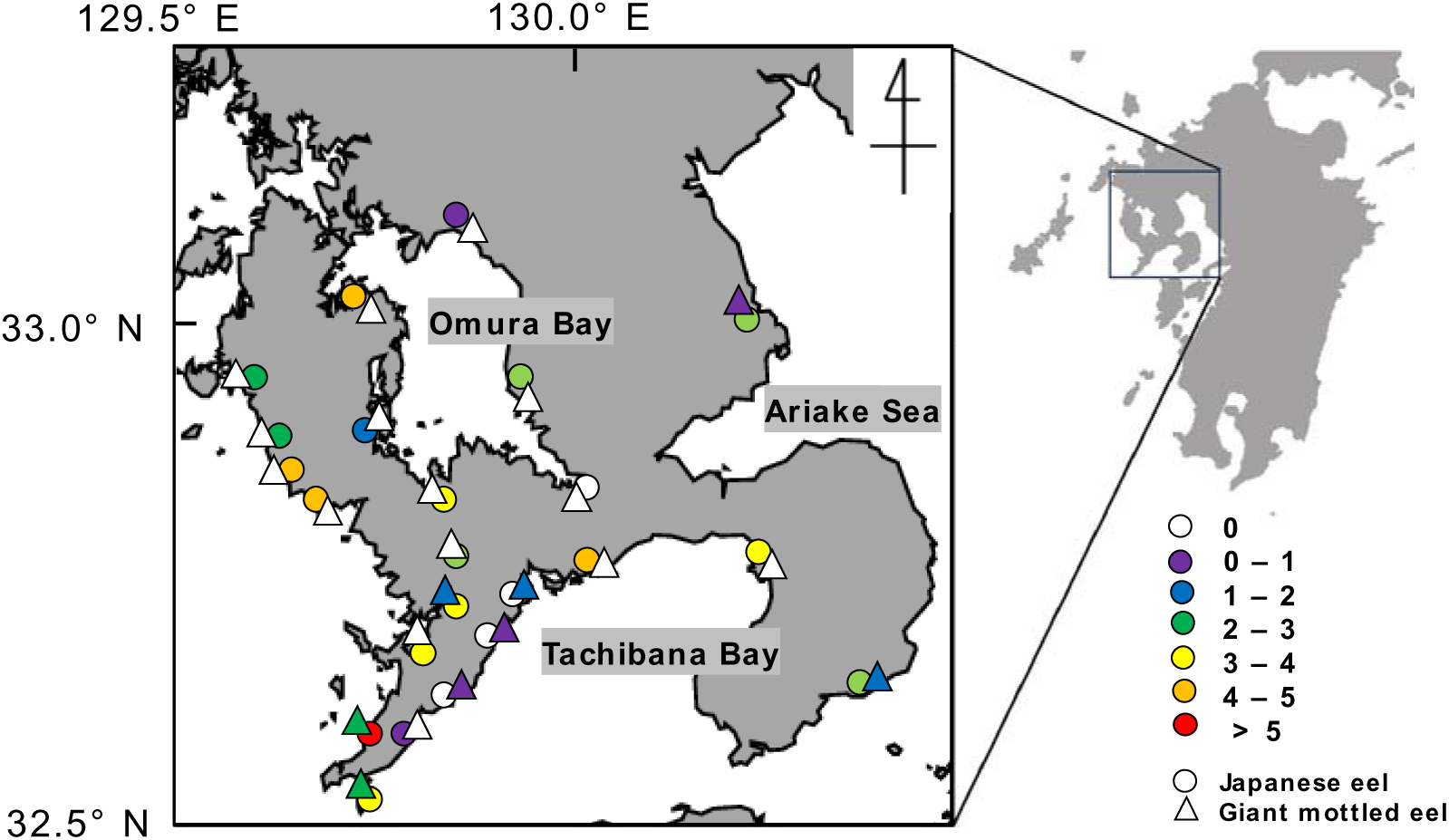
Map showing the geographic location of river mouths and environmental DNA (eDNA) concentrations for Japanese eels (circles) and giant mottled eels (triangles) in western Kyushu, Japan. Colors indicate mean values of logarithmic eDNA concentrations (log copies/L).

Water sampling was performed according to methods in the Manual for Aquatic eDNA Studies and Experiments (2021) and Ono et al. (2023). Briefly, water sampling was conducted in the main channel at each site. At sites where the main channel was inaccessible, sampling was carried out from a bridge using a fishing rod and reel with a disposable cup. After collecting 1 L of water using the disposable cup and pouch, 1 mL of 10% benzalkonium chloride solution (final concentration of 0.01% v/v) was immediately added to the sample to inhibit eDNA degradation (Yamanaka et al., 2017). All sampling cups and pouches were changed at each location, and the same surveyor carried out water sampling throughout the entire study. Powder-free nitrile gloves were worn during water sampling to avoid contamination. A pouch containing 1 L of distilled water and 1 mL of 10% benzalkonium chloride solution was also prepared for each sample as a field blank and a laboratory blank. Samples and blanks were transported in ice-filled coolers to the Fish and Ships Laboratory at Nagasaki University, Japan. They were vacuum filtered through one or two 47-mm GF/F glass filters (pore size: 0.7 µm, GE Healthcare, Tokyo, Japan) within 48 h. Filters were wrapped in aluminum foil and stored at -20°C until eDNA extraction.

### DNA extraction and qPCR analysis

eDNA extraction employed the methods of Itakura et al. (2019) and Ono et al. (2023). Briefly, eDNA was extracted from the filter using a DNeasy Blood and Tissue Kit (Qiagen, Hilden, Germany) and recovery was performed using a spin column method. Blanks were processed using the same method. Recovered eDNA and blanks were stored at -20°C until qPCR (Applied Biosystems 7500 Fast real-time PCR).

qPCR assays were performed according to the protocols of Itakura et al. (2019, 2020) and Ono et al. (2023). Species-specific primers for Japanese eel and giant mottled eel amplify 16SrRNA (Itakura et al., 2019; 2020, Ono et al., 2023). To ensure quantification, a calibration curve was also prepared using a dilution series of artificially synthesized genes. The synthetic gene sequence for Japanese eel is according to Ono et al. (2023), and for giant mottled eel is 5’-ACCCTAACAAGGCTGATTCGTTAACCGATTCACACACACACACACATCTCTCTCTCCC CACTAAATGTTGGAGGACATAAATG AGCAGTTATCCTGACATCTCTCTAATGCTATTCCTAATTACAATAAACATACTAGGGACT ACTACCATACACACATTCACACCAACAACCCAACTATCCATAAACATAGGATTTGCAGT CCCAATATGACTCGCC ACCGTAATCGTCGGGTATACGAAATCA-3’. Dilution series were adjusted to 1×10¹, 1×10², 1×10³, 1×10^5^, 1×10^7^ and 1×10^9^ copies for both species. The coefficients of determination (*R^2^*) of calibration curves for Japanese and giant mottled eels were 0.94-1.00 and 0.96-1.00, respectively. In each round of qPCR, three replicates per sample and pure water as a negative control were analyzed. Copy numbers were calculated based on calibration curves for each round and Ct values for each sample. No eDNA was detected in field or laboratory blanks, nor in any negative controls.

### Data analysis

To determine whether eDNA concentrations differ between regions in western Kyushu, tests for differences in mean eDNA concentrations were conducted for Japanese and giant mottled eels. The region was divided into three areas, North, along Omura Bay and South, according to the Japan Meteorological Agency (JMA) regional classification (JMA, https://www.jma.go.jp). eDNA concentrations were log-transformed (Doi et al., 2017; Maruyam et al., 2018), and the non-parametric Kruskal-Wallis test was employed because a normal distribution could not be assumed. Spearman rank correlation was used to determine the correlation between eDNA concentrations of Japanese and giant mottled eels at sites inhabited by both species.

Generalized Linear Models (GLMs) were constructed to identify environmental factors affecting eDNA concentrations of both species. Log-transformed eDNA concentration (> 0) was used as the dependent variable, and explanatory variables were water temperature, dissolved oxygen, water depth, flow velocity, river width, salinity, elevation and distance from the river mouth. As variables need to be independent, variance inflation factors were used to examine multicollinearity between variables. As a result, salinity (VIF > 5) was excluded (Zuur et al., 2009) (Table S1). GLM analysis used Gaussian distribution for family and identity as the link function. The best model was assumed to have the lowest value of the Akaike Information Criterion (AIC) from all possible regressions and a model with ΔAIC < 2 was assumed to be a reasonable alternative to the best model (Burnham and Anderson, 2010; Kasai et al., 2021).

## RESULTS

eDNA of Japanese eels was detected in 19 of 23 rivers (82.6%) and 56 of 87 sites (64.3%), whereas that of giant mottled eels was identified in 8 rivers (34.8%) and 16 sites (18.4%) (Fig. 1 and Fig. S1). eDNA of both species was found at 10 sites in 5 rivers (Fig. 1 and Fig. S1). eDNA for Japanese eels was detected at 6 of 9 sites in the North (66.7%), 13 of 23 sites in Omura (56.5%) and 37 of 55 sites in the South (67.3%). In contrast, giant mottled eel eDNA were detected at 1 of 9 sites in the North (11.1%), no sites in Omura and 15 of 55 sites in the South (27.3%).

eDNA concentrations of Japanese eels ranged from 3 to 30333 copies/L, whereas those of giant mottled eels ranged from 2 to 158 copies/L (Table S2). Mean eDNA concentrations (± S.D.) of Japanese eels by region were 2438 ± 6193 in the North, 1640 ± 3083 at Omura and 15868 ± 33897 in the South, with no significant differences (n = 56, *p* = 0.14) (Fig. 2a). On the other hand, mean eDNA concentrations of giant mottled eels were 3 ± 8 in North, 0 in Omura and 11 ± 29 in South, with significant differences between regions (n = 16, *p* = 0.01).

**Fig. 2.**
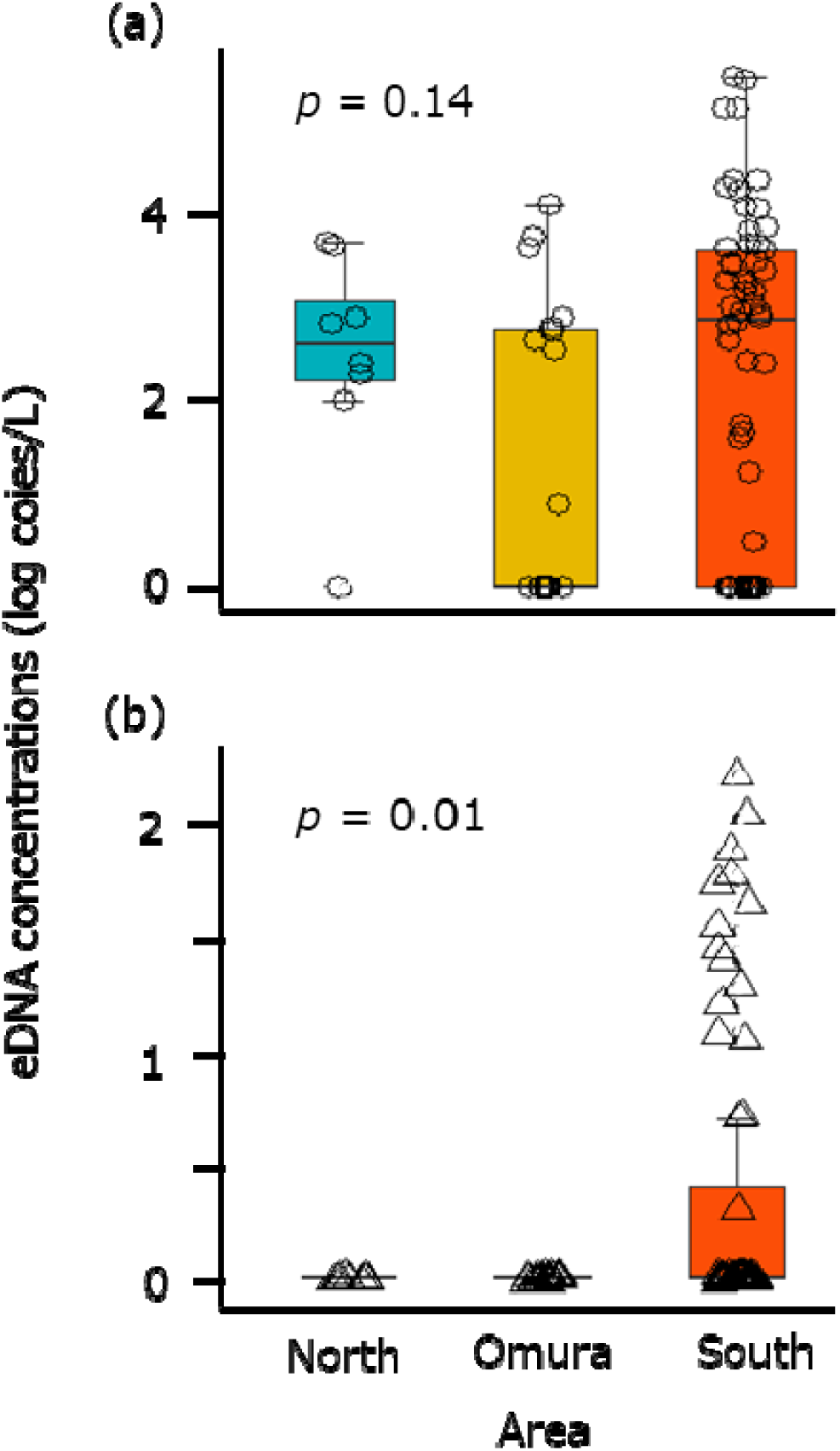
Comparison of environmental DNA concentrations by region in western Kyushu, Japan. (a) Japanese eel and (b) Giant mottled eel. Lower and upper box boundaries indicate the 25th and 75th percentiles. Horizontal lines represent the means. Lines inside boxes are the medians, and outer whiskers are the 10th and 90th percentiles.

Both Japanese and giant mottled eels were detected at 10 sites in 5 rivers (Figs. 1 and S1). There was no correlation between eDNA concentrations of the two species at the 10 sites where eDNA of both was detected (n = 10, *p* = 0.08) (Fig. 3). Japanese eels were more abundant than giant mottled eels at all 10 sites, but the lack of correlation suggests that they may adopt different microhabitats.

**Fig. 3.**
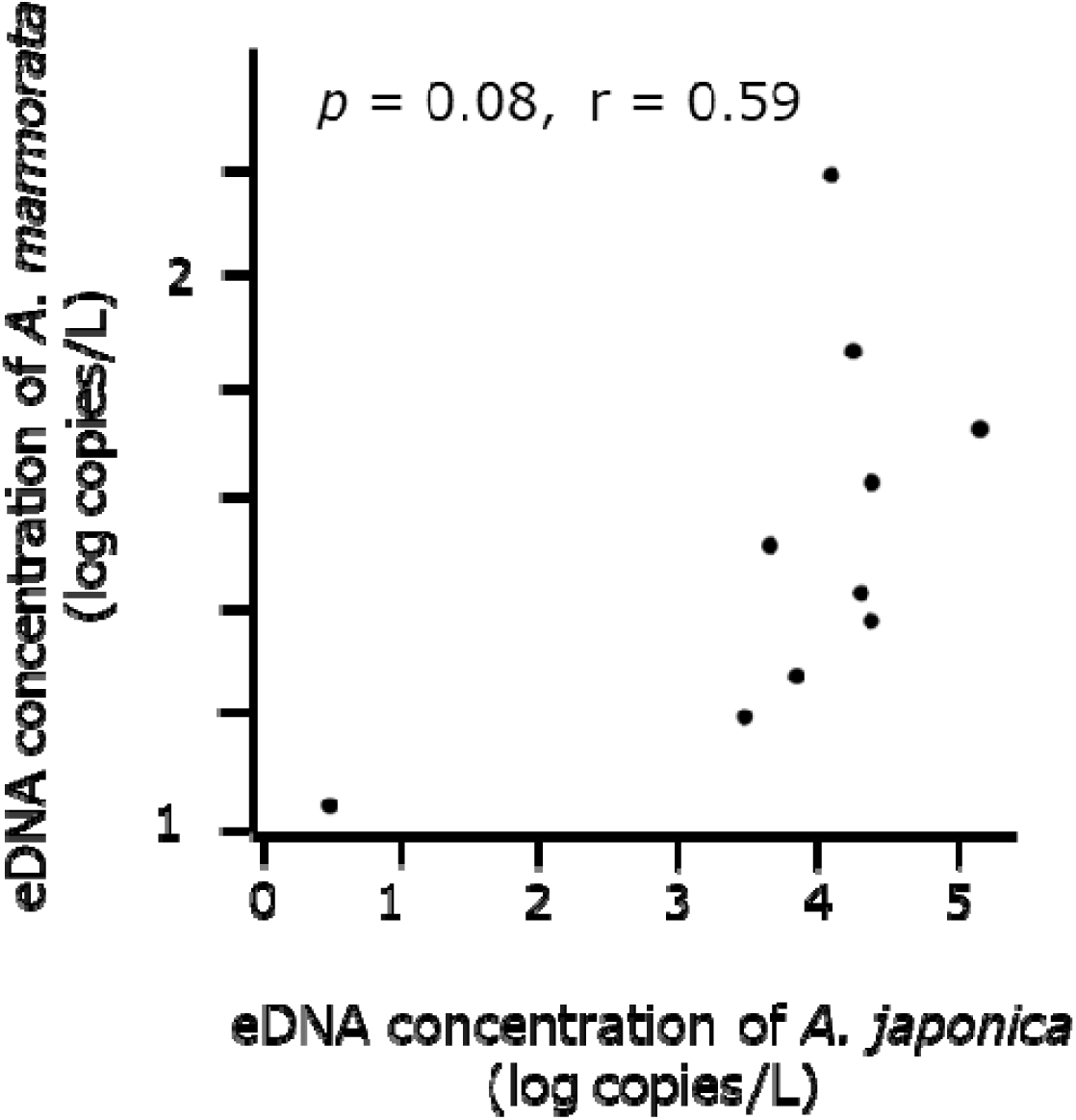
Relationship between environmental DNA concentrations at sites where both Japanese and giant mottled eels were detected in western Kyushu, Japan.

The best GLM model for Japanese eels included depth, velocity, temperature, dissolved oxygen and elevation, and for giant mottled eels, distance from the river mouth, river width and dissolved oxygen (Table 1). Within the range of ΔAIC < 2, there were eleven models for Japanese eels and ten models for giant mottled eels, and the direction of negative and positive effects within each species was consistent across all parameters (Table S3).

**Table 1.**
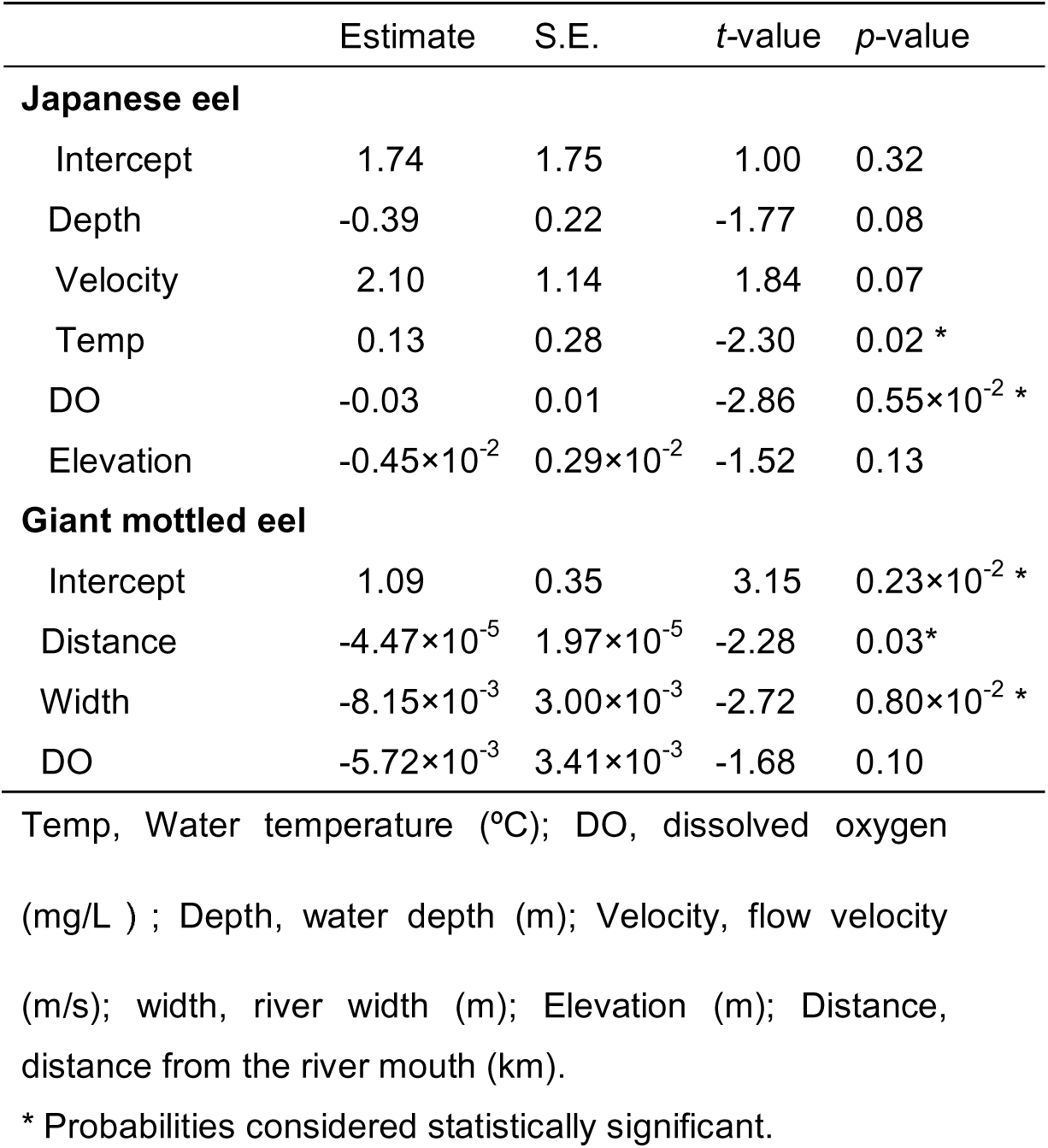
Summary of the best model by GLMs.

## DISCUSSION

Using eDNA methods, this study showed that there are distinct geographic distributional differences between Japanese and giant mottled eels along the northern west coast of Kyushu, Japan, supporting our hypothesis that distribution patterns of the two eel species differ within and between rivers, despite the limited geographic range. In Kyushu, the dominant *Anguilla* species is the Japanese eel (Kano et al., 2017), as confirmed by results of this study. We found that Japanese eels occur in nearly all rivers in all regions. On the other hand, giant mottled eel eDNA was not detected in rivers around Omura Bay, a closed inner bay, and it was also detected at low levels at northern sites (Figs. 1 and 2), supporting our hypothesis that abundance of the two species would differ among rivers. In addition, no correlation was found between the eDNA concentrations of the two species in 7 rivers inhabited by both species (Fig. 3).

Japanese eels are probably ubiquitous in rivers of the Japanese archipelago. Japanese eel eDNA was detected in most (82.6%) rivers examined in this study (Figs. 1 and 2a). Japanese eel eDNA was detected in rivers across the country, with the exception of the northern part of the Sea of Japan, in an extensive eDNA survey by Kasai et al. (2021). Ono et al. (2023) also reported detection in three small rivers in northwest Kyushu, consistent with this study’s high detection rates in all regions (Figs. 1 and 2). In addition, Japanese eels were detected in rivers flowing into Omura and Tachibana Bays (Fig. 1). In fact, ishi-kura fishing nets, which are underwater piles to catch wild dwelling eels (Kanzaki et al., 2018) have been set up and used to catch adult Japanese eel in rivers that flow into Omura Bay (unpublished, MY). The wide geographic distribution of this species may reflect extensive larval dispersal, i.e., that larvae are transported westward from their spawning grounds by the North Equatorial Current, reaching the western edge of the ocean basin near the Philippines, before moving northward to reach the wider East Asian region (Kuroki et al., 2014; Zenimoto et al., 2009).

Giant mottled eels are found in northwestern Kyushu, Japan, which may be the northern limit of their distribution. Based on previous reports (Kai 2009; Kai et al., 2011; 2012; 2018), we detected giant mottled eel eDNA in five rivers where they were previously unknown (Igayado, Arie, Tara, Otao, Chiji Rivers). Although giant mottled eels spawn in the same areas as Japanese eels and their larvae are also influenced by the Kuroshio and Tsushima Warm Currents to migrate up rivers (Kuroki et al., 2009; Mizuno & Nagasawa, 2009; Ogata *et al*., 2017). Giant mottled eels were not detected in rivers that flow into Omura Bay (Figs. 1 and 2b). Reasons for this are currently unknown, but we speculate that the extremely closed inner bay of Omura Bay makes it less susceptible to warm currents and that leptocephalus numbers of giant mottled eels are far smaller than those of Japanese eels (Kuroki et al., 2009). In addition, the detection rate was lower in the north than in the south (Fig. 2b). Kuroki et al (2006) claimed that for tropical species, including giant mottled eels, larval growth is faster and maximum larval sizes are smaller than for template eels such as the Japanese eel. Thus, southern rivers may constitute better habitat for giant mottled eels. In the future, it needs to be clarified whether giant mottled eels simply reach these rivers in smaller numbers, or whether settlement and growth limit their distribution.

Microhabitats in rivers where both species occur may not coincide. In this study, both species were detected at 10 sites in five rivers (Figs. 1 and S1). No correlation was found between eDNA concentrations of the two species at these sites, suggesting that their habitat preferences differ. The wide distribution of Japanese eels from freshwater to brackish water was also reported by Ono et al. (2023). In contrast, giant mottled eels were detected at shallower depths, around larger rocks in streams surrounded by thick woodlands. Giant mottled eels are reported to inhabit upstream areas where springs are present, while Japanese eels are commonly found downstream (Williamson et al., 1993; Ishibashi et al., 2001). Recent studies reported that giant mottled eels are found in midstream areas that are not adjacent to rice paddies, whereas Japanese eels are dominant in more downstream areas (Matsushige et al., 2022). Shiao et al. (2003) also found that giant mottled eels preferred deep pools in the river, and speculated that both interspecific competition and adaptive radiation occur. These suggest that Japanese eels inhabit diverse river environments from upstream to downstream, whereas giant mottled eels require complex river structures, such as deep pools in which to hide.

Additional studies will be required to identify preferred microhabitats of these species using eDNA concentrations. In the present study, there was a negative relationship between eDNA of Japanese eels and distance from the river mouth (Table 1 and S3). This result agreed with findings of Kasai et al. (2021). On the other hand, Japanese eel eDNA showed a negative correlation with depth, which conflicts with the results of Ono et al. (2023). For giant mottled eels, smaller rivers and higher elevations yielded higher eDNA concentrations (Table 1 and S3). This is consistent with the report by Matsushige et al. (2022) that giant mottled eels are found in midstream areas. Better results could be achieved by improving eDNA sampling methods, such as sampling multiple sites at the same location.

Limitations of this study include the lack of direct collection with electric shockers or eel pots, which prevented quantification of abundance and body size. However, Itakura et al. (2019) showed a significant weak positive correlation between the number of individuals directly captured and eDNA concentration. Ono et al. (2023) also reported higher concentrations at sites where Japanese eels were observed during eDNA collection. Furthermore, a large giant mottled eel was actually found in the Otao River, where the detection of eDNA was confirmed (personal communication, MY) (Fig. S1). Hence, eDNA concentrations in this study probably reflect eel biomass. As the abundance of both species has declined significantly (Mizuno, 2003), there is an urgent need to understand their dynamics, and eDNA techniques may be an effective, non-destructive tool (Doi et al., 2017; Jerde et al., 2013). In future studies, it will be important to determine annual changes in migration ecology and eDNA, in addition to metabarcoding analyses to understand interactions with other species.

Finally, juvenile eels are released by prefectural inland water fisheries co-operatives throughout Japan, except for Hokkaido and Nagasaki, to increase their populations; however, Wakiya et al. (2022) pointed out that releasing farmed eels does not necessarily increase populations, as released eels may have a negative impact on post-release survival and growth due to reduced capacity for intraspecific competition. It has also been claimed that releases can spread pathogens that are harmful to wild eels, and can significantly disrupt the balance of existing ecosystems (Miyai et al., 2004). Given these problems, cultured eels should not be released unnecessarily.

## Acknowledgements

This work was partially supported by JSPS KAKENHI (Grant Number JP21K06337 to M.Y.). We are grateful to students from the Faculty of Fisheries, Nagasaki University, for supporting field studies. Finally, we thank the editors and anonymous reviewers for their valuable comments and suggestions that greatly improved the quality of the manuscript.

## Supporting information

**Fig. S1.**
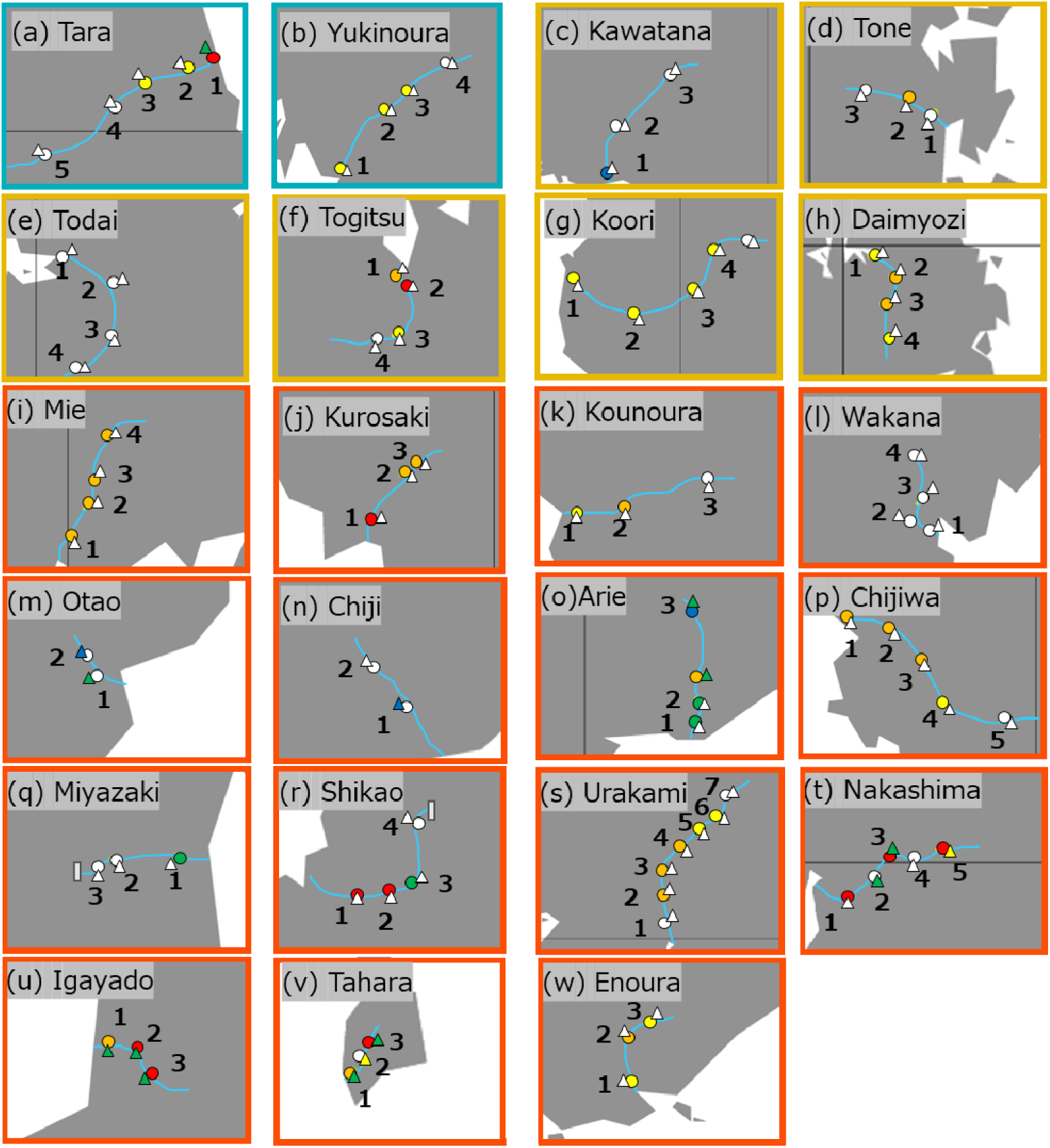
Map showing the geographic location of sampling sites and eDNA concentrations for Japanese eels (circles) and giant mottled eels (triangles) in western Kyushu, Japan. Rivers (a) and (b) are in the North. Those shown in (c)∼(h) are in Omura Bay and rivers (i)∼(w) are in the South. Numbers in each river are given in descending order of distance from the river mouth. Colors indicate values of logarithmic eDNA concentrations (log copies/L).

**Table S1.** Summary of variance inflation factor (VIF) analyses.

| Factor | Temp | DO | Depth | Velocity | Width | Salinity | Elevation | Distance |
| --- | --- | --- | --- | --- | --- | --- | --- | --- |
| VIF | 2.57 | 3.02 | 1.64 | 1.19 | 2.12 | 5.50 | 2.61 | 2.24 |

Temp, Water temperature (°C); DO, dissolved oxygen (mg/L ) ; Depth, water depth (m); Velocity, flow velocity (m/s); width, river width (m); Salinity (PSU); Elevation (m); Distance, distance from the river mouth (km)

**Table S2.** Environmental characteristics and concentrations of eDNA of Japanese eel (*Anguilla japonica*) and giant mottled eel (*Anguilla marmorata*) in rivers sampled in Nagasaki, Japan

| River | Site | Elevation (m) | Width (m) | Depth (m) | Water temp. (°C) | Velocity (m/s) | DO (%) | Salinity (PSU) | Distance from the river mouth (m) | eDNA of *A. japonica* (copies/L) | eDNA of *A. marmorata* (copies/L) |
| --- | --- | --- | --- | --- | --- | --- | --- | --- | --- | --- | --- |
| Urakami | S1 | 2 | 75 | 6.4 | 25.5 | 0.20 | 76.9 | 33.17 | 1000 | - | - |
|  | S2 | 7 | 4 | 3.8 | 26.1 | 0.29 | 46.8 | 31.58 | 2200 | 1033 | - |
|  | S3 | 2 | 11 | 0.2 | 31.2 | 0.16 | 150.1 | 0.10 | 3660 | 1102 | - |
|  | S4 | 8 | 8 | 0.1 | 30.6 | 0.30 | 123.7 | 0.08 | 4740 | 7312 | - |
|  | S5 | 34 | 5 | 0.1 | 29.3 | 0.17 | 130.8 | 0.11 | 6470 | 266 | - |
|  | S6 | 58 | 3 | 0.2 | 26.5 | 0.40 | 114.8 | 0.11 | 7230 | 258 | - |
|  | S7 | 95 | 6 | 0.1 | 25.4 | 0.29 | 110.5 | 0.11 | 8600 | - | - |
| Nakashima | S1 | 5 | 3 | 2.8 | 26.0 | 0.01 | 55.6 | 32.25 | 570 | 142702 | - |
|  | S2 | 7 | 2 | 1.7 | 24.8 | 0.01 | 102.7 | 0.07 | 1310 | - | 59 |
|  | S3 | 9 | 6 | 0.2 | 29.8 | 0.48 | 104.5 | 0.05 | 1810 | 23551 | 24 |
|  | S4 | 22 | 7 | 0.1 | 27.6 | 0.40 | 108.3 | 0.05 | 2700 | - | - |
|  | S5 | 36 | 7 | 0.3 | 26.6 | 0.26 | 100.7 | 0.05 | 3250 | 12188 | 158 |
| Togitsu | S1 | 1 | 21 | 0.1 | 30.0 | 0.10 | 90.3 | 25.94 | 500 | 5956 | - |
|  | S2 | 3 | 7 | 0.5 | 28.5 | 0.19 | 127 | 0.14 | 860 | 13200 | - |
|  | S3 | 30 | 5 | 0.1 | 23.1 | 0.08 | 96.8 | 0.13 | 2100 | 631 | - |
|  | S4 | 56 | 4 | 0.7 | 25.6 | 0.07 | 128.1 | 0.14 | 2800 | - | - |
| Tone | S1 | 1 | 22 | 1.3 | 29.8 | 0.07 | 68.9 | 32.57 | 530 | - | - |
|  | S2 | 4 | 44 | 0.3 | 24.2 | 0.27 | 99.9 | 0.04 | 1400 | 4467 | - |
|  | S3 | 47 | 10 | 0.3 | 21.4 | 0.06 | 101.2 | 0.03 | 3000 | - | - |
| Daimyozi | S1 | 0 | 72 | 1.6 | 28.1 | 0.33 | 97.8 | 33.28 | 440 | 672 | - |
|  | S2 | 5 | 32 | 1.4 | 29.5 | 0.10 | 73.6 | 32.95 | 1800 | 4619 | - |
|  | S3 | 15 | 6 | 0.1 | 25.9 | 0.03 | 91.9 | 0.04 | 3300 | 5175 | - |
|  | S4 | 20 | 3 | 0.1 | 23.6 | 0.04 | 112.7 | 0.04 | 5300 | 781 | - |
| Tahara | S1 | 3 | 5 | 0.2 | 25.9 | 0.06 | 72.6 | 0.12 | 0 | 6855 | 19 |
|  | S2 | 10 | 2 | 0.1 | 23.0 | 0.06 | 91.5 | 0.10 | 410 | - | 103 |
|  | S3 | 24 | 1 | 0.1 | 21.0 | 0.09 | 90.4 | 0.10 | 800 | 17525 | 75 |
| Igayado | S1 | 3 | 3 | 0.2 | 25.9 | 0.12 | 72.6 | 25.66 | 0 | 4419 | 33 |
|  | S2 | 10 | 6 | 0.1 | 23.0 | 0.10 | 91.5 | 0.14 | 390 | 23807 | 43 |
|  | S3 | 24 | 3 | 0.1 | 21.0 | 0.10 | 90.4 | 0.12 | 900 | 142702 | 54 |
| Shikao | S1 | 12 | 8 | 0.3 | 25.8 | 0.20 | 100.1 | 1.96 | 1500 | 303333 | - |
|  | S2 | 11 | 8 | 0.7 | 26.5 | 0.15 | 101.9 | 0.05 | 2450 | 285270 | - |
|  | S3 | 32 | 22 | 1.4 | 26.2 | 0.10 | 101.7 | 0.05 | 3100 | 39 | - |
|  | S4 | 32 | 3 | 0.1 | 30.8 | 0.38 | 140.8 | 0.07 | 4500 | - | - |
| Mie | S1 | 5 | 18 | 0.7 | 29.6 | 0.06 | 82.6 | 31.22 | 310 | 1848 | - |
|  | S2 | 6 | 5 | 0.4 | 23.2 | 0.17 | 90.8 | 0.46 | 740 | 2006 | - |
|  | S3 | 7 | 4 | 0.1 | 23.1 | 0.15 | 100.8 | 0.46 | 1160 | 2022 | - |
|  | S4 | 28 | 7 | 0.1 | 23.9 | 0.25 | 99.5 | 0.43 | 2000 | 1531 | - |
| Kurosaki | S1 | 5 | 22 | 0.2 | 22.3 | 0.50 | 105.8 | 0.05 | 220 | 11692 | - |
|  | S2 | 37 | 4 | 0.2 | 21.3 | 0.45 | 101.9 | 0.04 | 2000 | 4737 | - |
|  | S3 | 67 | 13 | 0.2 | 21.4 | 0.24 | 102.0 | 0.04 | 2500 | 4153 | - |
| Kounoura | S1 | 5 | 34 | 1.1 | 28.0 | 0.10 | 88.7 | 29.73 | 820 | 843 | - |
|  | S2 | 38 | 22 | 0.6 | 24.6 | 0.26 | 104.4 | 0.03 | 2700 | 1493 | - |
|  | S3 | 138 | 10 | 0.6 | 20.4 | 0.28 | 102.9 | 0.03 | 5500 | - | - |
| Yukinoura | S1 | 0 | 80 | 1.0 | 28.9 | 0.05 | 100.6 | 21.58 | 460 | 256 | - |
|  | S2 | 4 | 25 | 1.0 | 25.2 | 0.25 | 106.2 | 0.30 | 3800 | 100 | - |
|  | S3 | 10 | 15 | 0.9 | 24.8 | 0.47 | 104.9 | 0.02 | 5000 | 197 | - |
|  | S4 | 60 | 16 | 1.0 | 22.1 | 0.43 | 98.4 | 0.03 | 8000 | - | - |
| Arie | S1 | 7 | 6 | 0.1 | 27.0 | 0.24 | 119.8 | 0.07 | 400 | 45 | - |
|  | S2 | 20 | 13 | 0.3 | 26.0 | 0.22 | 112.0 | 0.06 | 1500 | 17 | - |
|  | S3 | 38 | 12 | 0.1 | 19.7 | 0.17 | 104.9 | 0.04 | 2500 | 2906 | 16 |
|  | S4 | 190 | 8 | 0.1 | 27.1 | 0.23 | 111.0 | 0.07 | 6300 | 3 | 11 |
| Tidiwa | S1 | 0 | 10 | 0.2 | 27.4 | 0.54 | 116. | 0.09 | 620 | 4895 | - |
|  | S2 | 29 | 4 | 0.1 | 22.8 | 0.76 | 4 | 0.04 | 2400 | 3156 | - |
|  | S3 | 107 | 6 | 0.4 | 22.5 | 0.16 | 104. | 0.04 | 3900 | 2483 | - |
|  | S4 | 153 | 8 | 0.1 | 19.6 | 0.69 | 7 | 0.04 | 5100 | 699 | - |
|  | S5 | 292 | 2 | 0.3 | 18.7 | 0.41 | 103. | 0.05 | 7500 | - | - |
|  |  |  |  |  |  |  | 104.<br>8 |  |  |  |  |
|  |  |  |  |  |  |  | 103.<br>8 |  |  |  |  |
| Todai | S1 | 0 | 102 | 0.6 | 31.2 | 0.14 | 115. | 30.78 | 0 | - | - |
|  | S2 | 2 | 30 | 0.8 | 27.5 | 0.08 | 3 | 0.05 | 2700 | - | - |
|  | S3 | 27 | 9 | 0.1 | 28.8 | 0.37 | 125. | 0.06 | 5500 | - | - |
|  | S4 | 67 | 2 | 0.3 | 24.8 | 0.12 | 6 | 0.04 | 7800 | - | - |
|  |  |  |  |  |  |  | 134.<br>6 |  |  |  |  |
|  |  |  |  |  |  |  | 97.4 |  |  |  |  |
| Koori | S1 | 0 | 58 | 1.1 | 30.3 | 0.03 | 63.6 | 30.52 | 320 | 781 | - |
|  | S2 | 44 | 11 | 1.2 | 25.2 | 0.05 | 111.2 | 0.04 | 5300 | 443 | - |
|  | S3 | 110 | 11 | 0.1 | 23.1 | 0.16 | 105.7 | 0.03 | 9300 | 576 | - |
|  | S4 | 178 | 10 | 0.1 | 21.3 | 0.56 | 100.3 | 0.03 | 12000 | 348 | - |
|  | S5 | 249 | 11 | 0.2 | 20.8 | 0.45 | 104.2 | 0.03 | 14000 | - | - |
| Kawatana | S1 | 1 | 92 | 1.7 | 29.6 | 0.10 | 89.5 | 31.81 | 560 | 78 | - |
|  | S2 | 16 | 22 | 0.6 | 27.6 | 0.10 | 128.3 | 0.08 | 6000 | - | - |
|  | S3 | 59 | 11 | 0.4 | 24.2 | 0.0025 | 95.5 | 0.05 | 15600 | - | - |
| Tara | S1 | 3 | 32 | 0.2 | 28.6 | 0.05 | 68.6 | 25.77 | 220 | 19939 | 27 |
|  | S2 | 16 | 16 | 0.2 | 27.1 | 0.11 | 108.4 | 0.04 | 1500 | 795 | - |
|  | S3 | 54 | 14 | 0.1 | 24.4 | 0.46 | 111.9 | 0.03 | 3500 | 657 | - |
|  | S4 | 113 | 8 | 0.1 | 21.6 | 0.26 | 106.9 | 0.03 | 5500 | - | - |
|  | S5 | 434 | 5 | 0.4 | 18.4 | 0.23 | 104.4 | 0.02 | 10100 | - | - |
| Miyazaki | S1 | 2 | 12 | 0.1 | 23.5 | 0.15 | 68.9 | 19.14 | 410 | 55 | - |
|  | S2 | 8 | 5 | 0.2 | 21.4 | 0.10 | 100.5 | 0.09 | 1300 | - | - |
|  | S3 | 7 | 7 | 0.1 | 22.9 | 0.25 | 98.8 | 0.07 | 1600 | - | - |
| Chiji | S1 | 4 | 7 | 0.2 | 21.7 | 0.35 | 94.0 | 0.05 | 160 | - | 2 |
|  | S2 | 34 | 11 | 0.2 | 21.5 | 0.13 | 98.4 | 0.04 | 770 | - | - |
| Wakana | S1 | 3 | 20 | 1.2 | 26.2 | 0.10 | 95.8 | 31.40 | 90 | - | - |
|  | S2 | 8 | 5 | 0.2 | 20.4 | 0.35 | 103.2 | 0.08 | 1650 | - | - |
|  | S3 | 14 | 10 | 0.1 | 22.7 | 0.11 | 105. | 0.08 | 2150 | - | - |
|  |  |  |  |  |  |  | 5 |  |  |  |  |
|  | S4 | 52 | 4 | 0.1 | 21.1 | 0.50 | 100.6 | 0.02 | 3600 | - | 5 |
| Otao | S1 | 5 | 8 | 0.2 | 21.4 | 0.08 | 101.6 | 0.06 | 132 | - | 12 |
|  | S2 | 21 | 4 | 0.1 | 22.4 | 0.04 | 106.0 | 0.06 | 430 | - | 5 |
| Enoura | S1 | 0 | 49 | 0.8 | 25.0 | 0.09 | 91.1 | 23.11 | 1100 | 892 | - |
|  | S2 | 4 | 5 | 0.2 | 24.8 | 0.11 | 110.9 | 0.13 | 2500 | 3090 | - |
|  | S3 | 26 | 4 | 0.1 | 22.4 | 0.18 | 97.4 | 0.12 | 3230 | 445 | - |

**Table S3.**
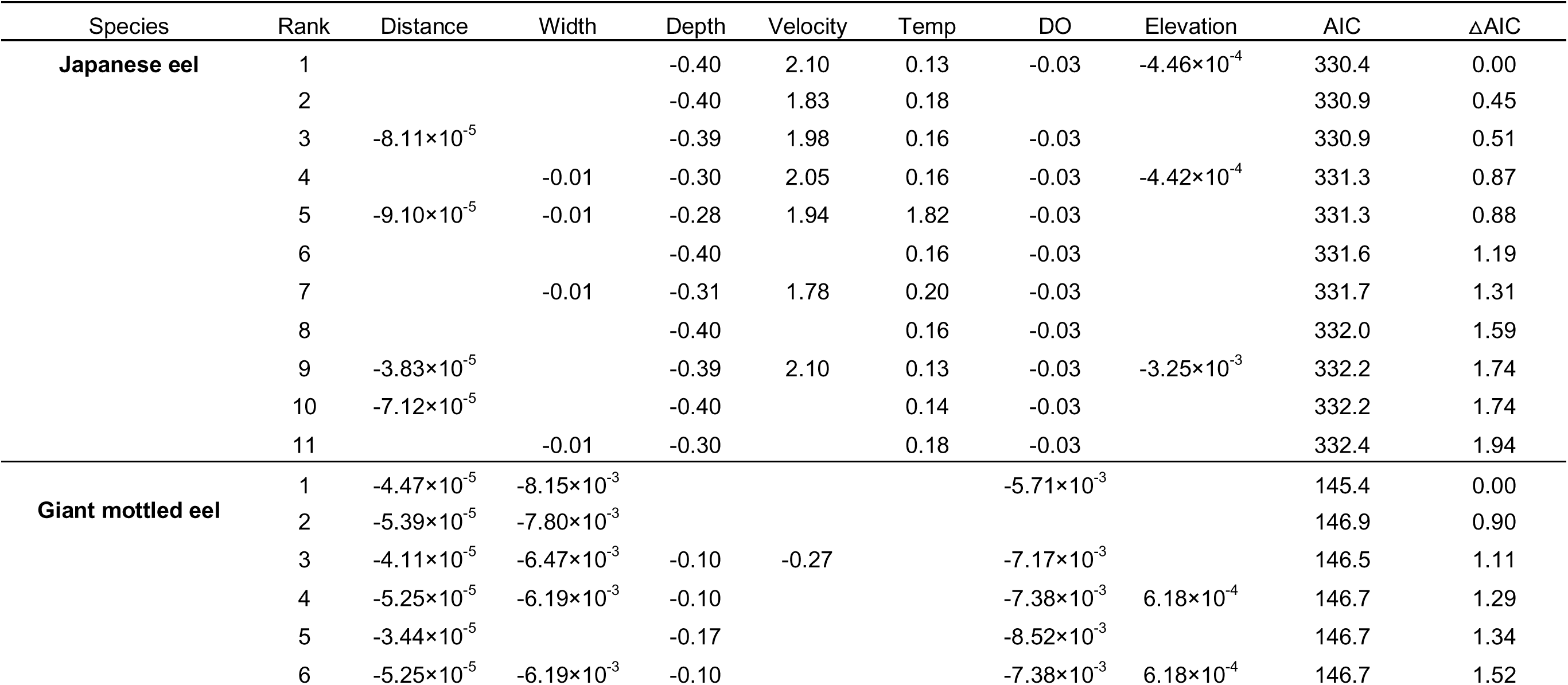
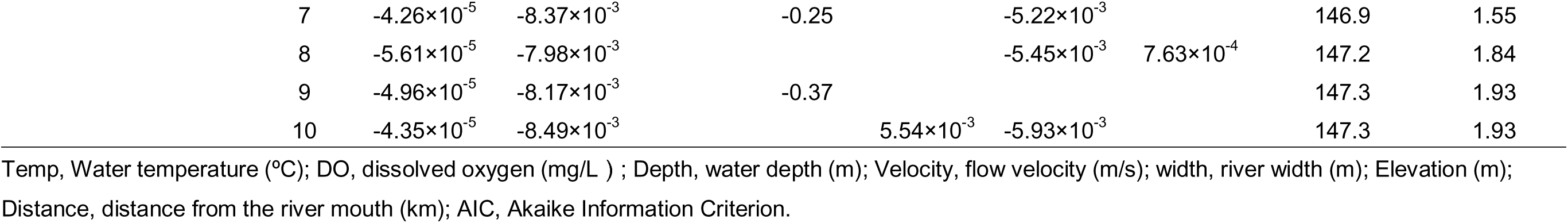
Summary of GLMs with ΔAIC < 2.

